# Cardiomyocytes possess an intrinsic catecholaminergic machinery that regulates cellular homeostasis and electrophysiological stability

**DOI:** 10.64898/2026.06.29.735427

**Authors:** Dimitra Krexi, Daniele Linardi, Charles Redwood

## Abstract

**Background:** Catecholamines play a central role in cardiac performance, coordinating myocardial contractility, conduction, metabolism, and electrophysiological stability. In the heart, their actions have been attributed primarily to sympathetic nerve terminals and circulating adrenal catecholamines. The discovery of an intrinsic non-neuronal cholinergic system within cardiomyocytes challenges this neurocentric paradigm and raises the possibility that cardiomyocytes also possess an intrinsic catecholaminergic programme. Here, we investigated whether cardiomyocytes possess an intrinsic catecholaminergic programme and its contribution to cardiomyocyte homeostasis and stress responses.

**Methods:** We investigated catecholamine biosynthesis and handling in human induced pluripotent stem cell-derived cardiomyocytes, adult mouse cardiomyocytes, H9C2 cells, rat ventricular tissue, and Langendorff-perfused mouse hearts. Protein expression of catecholamine biosynthetic enzymes and vesicular monoamine transporters was assessed by immunoblotting and immunohistochemistry, while vesicular monoamine uptake was evaluated using fluorescent false neurotransmitters. Functional consequences of catecholamine biosynthesis inhibition were examined using pharmacological approaches, assessing cell viability, apoptosis, organelle homeostasis, metabolic signalling, and cardiac electrophysiology.

**Results:** Tyrosine hydroxylase, aromatic L-amino acid decarboxylase, dopamine β-hydroxylase, and vesicular monoamine transporters were detected in cardiomyocytes across models. Expression of catecholamine biosynthetic enzymes increased following ischaemia–reperfusion injury in rat heart tissue (TH p=0.008, AADC p=0.031, DBH p=0.008). Pharmacological inhibition of catecholamine biosynthesis caused dose-dependent reductions in cardiomyocyte viability (p<0.0001), increased apoptosis, organelle stress, and mitochondrial dysfunction, with greater effects under oxidative stress. Mechanistically, catecholamine depletion suppressed mTORC1 signalling and activated LKB1–AMPK–ULK1 pathways. In Langendorff-perfused hearts, tyrosine hydroxylase inhibition induced ventricular arrhythmias in 5 of 6 hearts, including sustained ventricular tachycardia, polymorphic ventricular tachycardia, and ventricular fibrillation.

**Conclusions:** These findings identify cardiomyocytes as previously unrecognised catecholamine-competent cells expressing intrinsic machinery for catecholamine biosynthesis and vesicular handling. Disruption of this pathway compromises metabolic and organelle homeostasis, activates energy-stress and autophagy-related signalling, and promotes malignant ventricular arrhythmias. Intrinsic cardiomyocyte catecholamine biology therefore represents a non-neuronal regulatory axis essential for myocardial resilience and electrical stability, with potential relevance to ischaemic injury and stress-induced dysfunction.

## 1. INTRODUCTION

Catecholamines are central regulators of cardiac function, modulating heart rate, contractility, conduction, and electrophysiological stability through sympathetic innervation and adrenergic receptor signalling (1-3). Perturbations in catecholamine signalling contribute to myocardial injury, maladaptive remodelling, heart failure, and arrhythmogenesis (4-6). Beyond their canonical haemodynamic and electrophysiological actions, catecholamines also influence cardiomyocyte metabolism and cellular homeostasis, including modulation of mTOR signalling, and energy-sensing pathways (7).

Catecholamine biosynthesis occurs through a conserved enzymatic cascade in which tyrosine hydroxylase (TH) and aromatic L-amino acid decarboxylase (AADC) generate dopamine, dopamine β-hydroxylase (DBH) converts dopamine to norepinephrine, and phenylethanolamine N-methyltransferase (PNMT) catalyses the conversion of norepinephrine to epinephrine (8, 9). Following synthesis, monoamines are sequestered into secretory vesicles, released in response to stimulation, and subsequently cleared from the extracellular space via dopamine and norepinephrine transporters (DAT and NET respectively) (figure 1) (10-12).

**Figure 1.**
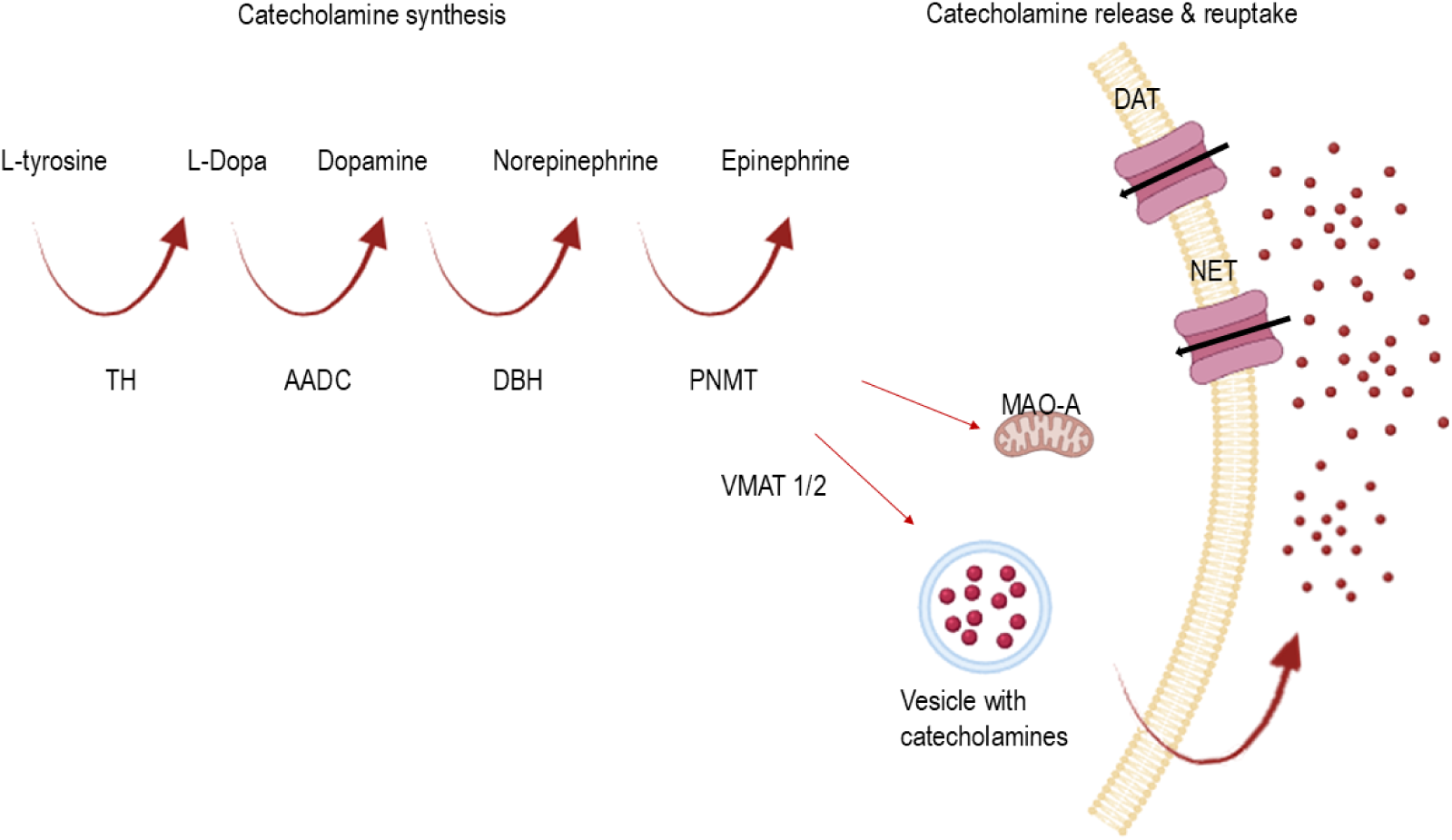
Catecholamines synthesis, release and reuptake pathway.

Within the heart, catecholamines are classically viewed as extrinsic mediators derived from sympathetic nerve terminals, which release norepinephrine locally within the myocardium, and from circulating sources such as the adrenal medulla (13, 14). However, this conventional model has been challenged by the recognition that cardiomyocytes possess intrinsic neurotransmitter-related systems, including a non-neuronal cholinergic network capable of modulating cardiac electrophysiology and stress responses (15). These findings raise the possibility that cardiomyocytes may also harbour intrinsic monoaminergic machinery with local regulatory functions.

Whether cardiomyocytes possess a functional catecholaminergic machinery capable of catecholamine synthesis, handling, and stress-responsive regulation remains unclear. Here, we identify an intrinsic catecholaminergic programme in cardiomyocytes and define its contribution to cellular homeostasis at baseline and in response to hydrogen peroxide (H₂O₂)-induced oxidative stress.

## 2. METHODS

### 2.1 Human induced pluripotent stem cell-derived cardiomyocytes, adult mouse cardiomyocytes, and H9C2 cells

Human induced pluripotent stem cells carrying a titin N-terminal enhanced green fluorescent protein tag were maintained in culture and differentiated into cardiomyocytes by Wnt pathway modulation. Cells were cultured in RPMI 1640/B27 medium minus insulin and maintained until spontaneous beating was established (16).

Adult ventricular cardiomyocytes were isolated from male CD1 mice by retrograde perfusion and enzymatic digestion (17). N=4.

H9C2 cardiomyoblasts were cultured in DMEM with 10% fetal bovine serum [FBS, PAA], 2 mM L-glutamine, and 0.5% penicillin and streptomycin; all Life Technologies). They were treated with three doses (25μM, 50 μM and 100 μM) of the tyrosine hydroxylase inhibitor, metyrosine (Selleckchem), three doses (25μM, 50 μM and 100 μM) of the aromatic L-amino acid decarboxylase inhibitor, carbidopa (Tocris) and three doses (5μM, 10 μM and 20 μM) of the nepicastat hydrochloride (Tocris) for 24 hours. Hydrogen peroxide stress was induced with 600μM H2O2 treatment for 6 hours. N=3 (18).

### 2.2 Fluorescent false neurotransmitter uptake assays (FFNs)

Catecholamine handling was assessed in H9C2 cells using the fluorescent false neurotransmitters FFN102 and FFN270. Cells were incubated with each probe and visualised by confocal microscopy using an Olympus FV3000. Nomifensine and reserpine were used as mechanistic controls to assess transporter-dependent uptake and vesicular retention (19)(20).

### 2.3 Myocardial ischemia-reperfusion (I/R) rat model

An adult Sprague-Dawley rat regional ischemia-reperfusion (I/R) injury model, through the left anterior descending (LAD) coronary ligation, was used to evaluate in vivo regulation of the catecholamine pathway (21). Following anaesthesia and ventilation support, thoracotomy was performed, and the LAD coronary artery was encircled with a suture and ligated for 30 min. The snare was then released for myocardial reperfusion. The operating procedure and post-surgical animal care were in accordance with European legislation and approved by local Italian authorities. After two weeks of recovery, the rats were sacrificed, their hearts were fixed in 4% paraformaldehyde solution for 48 hours for subsequent tissue section. As control, hearts from sham-operated (thoracotomy only), age-matched rats were taken.

### 2.4 Immunohistochemistry

Rat tissue was sectioned at 4μm thickness. Initially, tissue was deparaffinised and rehydrated with xylene and reducing percentage of ethanol. Antigen retrieval was achieved with boiling for 25 minutes. Rabbit specific HRP/DAB Detection IHC Kit by Abcam (Cat #ab64261) was used according to manufacturer’s protocol. Primary antibodies were diluted in 3% BSA and samples were incubated at 4°C in primary antibodies overnight. Primary antibodies are in detail in the online supplement. The immunoreactive score (IRS) of Remmele and Stegner score is used to measure the intensity of a cytoplasmic reaction and to assess the expression of proteins (22). An Evos M7000 microscope was used to visualize the tissue.

### 2.5 Immunocytochemistry

Isolated cardiomyocytes from adult wild type CD1 mice Cells were fixated with 4% paraformaldehyde for 15 minutes, followed by permeabilization. Cells were blocked with 10% goat serum and then were incubated with TH and AADC primary antibody at 4°C overnight. Alexa Fluor-conjugated secondary antibodies at 1:200 dilution was used for secondary immunostaining. Olympus FV3000 was used to visualise the cells.

### 2.6 Western Blotting

HiPSC-CMs, CD1 ventricular cardiomyocytes and H9C2 cells were frozen in RIPA buffer with protease and phosphatase inhibitors. Samples were diluted in LDA buffer. After heating them at 95°C for 5 minutes, samples were added to NuPAGE 12% Bis-Tris gel for electrophoresis. Wet transfer was performed followed by blocking with 1% BSA. Protein signals were detected using an ODYSSEY® M imaging system.

### 2.7 Cellular assays

Cell viability, cytotoxicity, apoptosis, reactive oxygen species production, mitochondrial membrane potential, lysosomal content, mitochondrial integrity, and endoplasmic reticulum stress were assessed using fluorescence- and plate-based assays (Online supplement).

### 2.8 Langendorff Perfused hearts

Langendorff-perfused heart experiments were performed in 6 young adult C57BL/6 wild type male mice. All mice were sacrificed by cervical dislocation according to Schedule 1 of the UK Home Office Animals Scientific Procedures Act (1986). After thoracotomy the hearts were excised and transferred immediately to cold Ca2+-free Krebs solution containing heparin. The hearts were cannulated to a 23 G needle via the aorta and subsequently mounted onto the Langendorff system. Two ECG leads were placed, one to the atrium and the other one to the apex on the opposite side.

To investigate the effect of TH inhibition in the heart, Krebs solutions containing the following solvents were perfused in a sequence: Solvent 1 (Krebs solution), Solvent 2 (50μM TH inhibitor), Solvent 3 (Krebs solution). All solutions were perfused at 3.4 ml/min with the temperature of water bath being 36 ± 1 °C. Solvent 2 and 3 were perfused for 30minutes. Burst pacing was also performed. Three short bursts of rapid pacing were applied each time starting from 90 ms and reducing by 10ms/cycle each time with 30ms as the final burst pacing.

### 2.12 Statistical analysis

Comparisons between two groups of normally distributed data are performed by t test and non-normally distributed data by Mann-Whitney. For multiple group comparisons Kruskal-Wallis or Wilcoxon are used. p value < 0.05 is considered as statistically significant. Statistical analysis is performed using SPSS version 29.

## 3. RESULTS

### 3.1 Cardiomyocytes express key elements of the catecholamine biosynthetic machinery across species and model systems

TH was detected in human induced pluripotent stem cell-derived cardiomyocytes (hiPSC-CMs), indicating the presence of the rate-limiting enzyme required for the conversion of tyrosine to L-DOPA. AADC, DBH, and PNMT were also detected, demonstrating the presence of the key enzymatic cascade in hiPSC-CMs required for catecholamine biosynthesis.

This expression profile was conserved across species and model systems, as TH, AADC, DBH, and PNMT expression was also confirmed in isolated ventricular cardiomyocytes from adult male wild-type CD1 mice and in the rat cardiomyoblast cell line, H9C2 (Figure 2).

**Figure 2.**
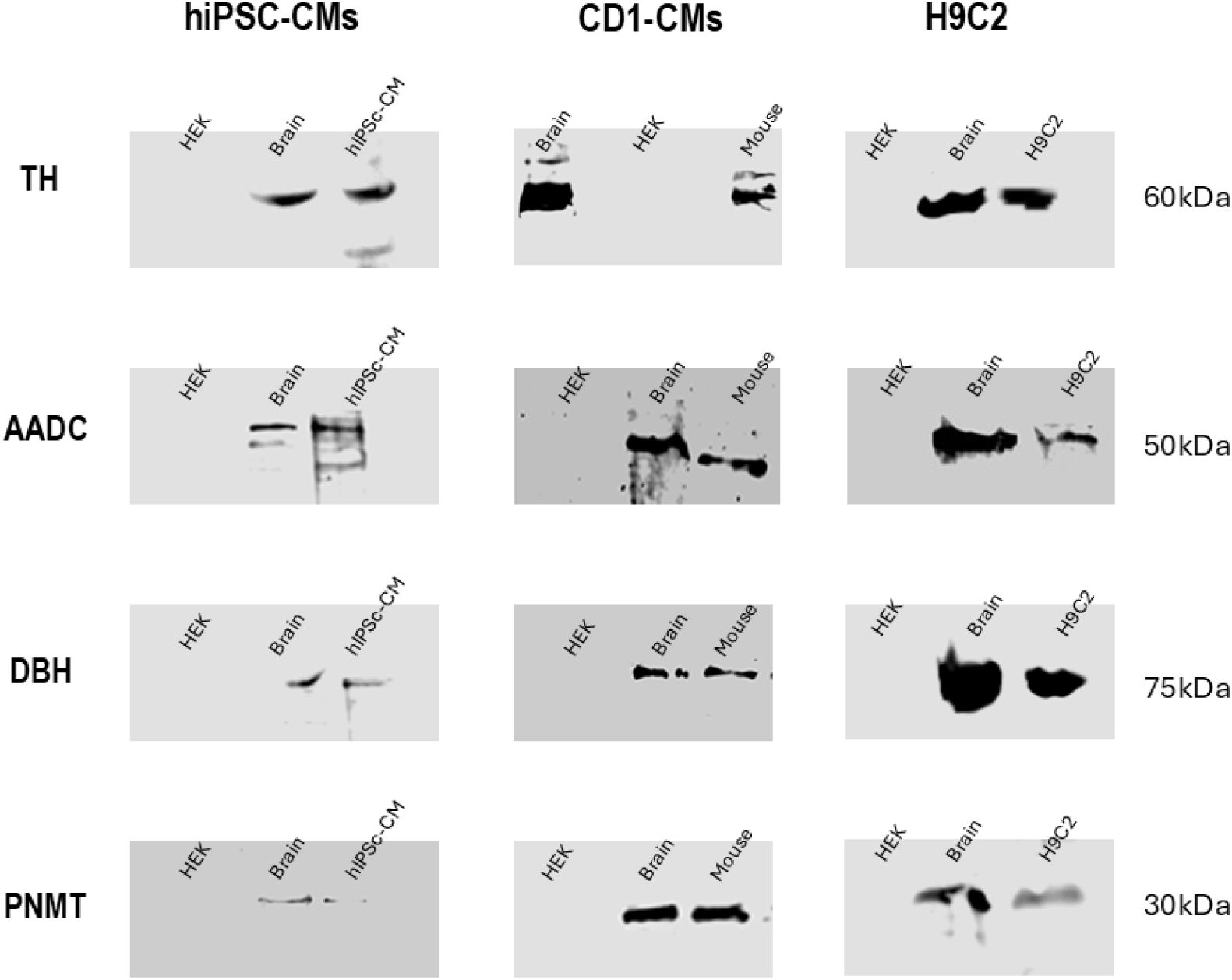
Cardiomyocytes express catecholamine biosynthetic enzymes across species and model systems. Representative western blots demonstrating expression of TH, AADC, DBH and PNMT in human induced pluripotent stem cell-derived cardiomyocytes (hiPSC-CMs), isolated adult CD1 mouse ventricular cardiomyocytes, and H9C2 cells. Mouse brain lysate was used as a positive control and HEK cells as negative control. Molecular weight markers (kDa) are shown.

TH and AADC exhibited predominantly cytosolic localisation, with diffuse intracellular staining observed throughout the cardiomyocytes (Figure 3).

**Figure 3.**
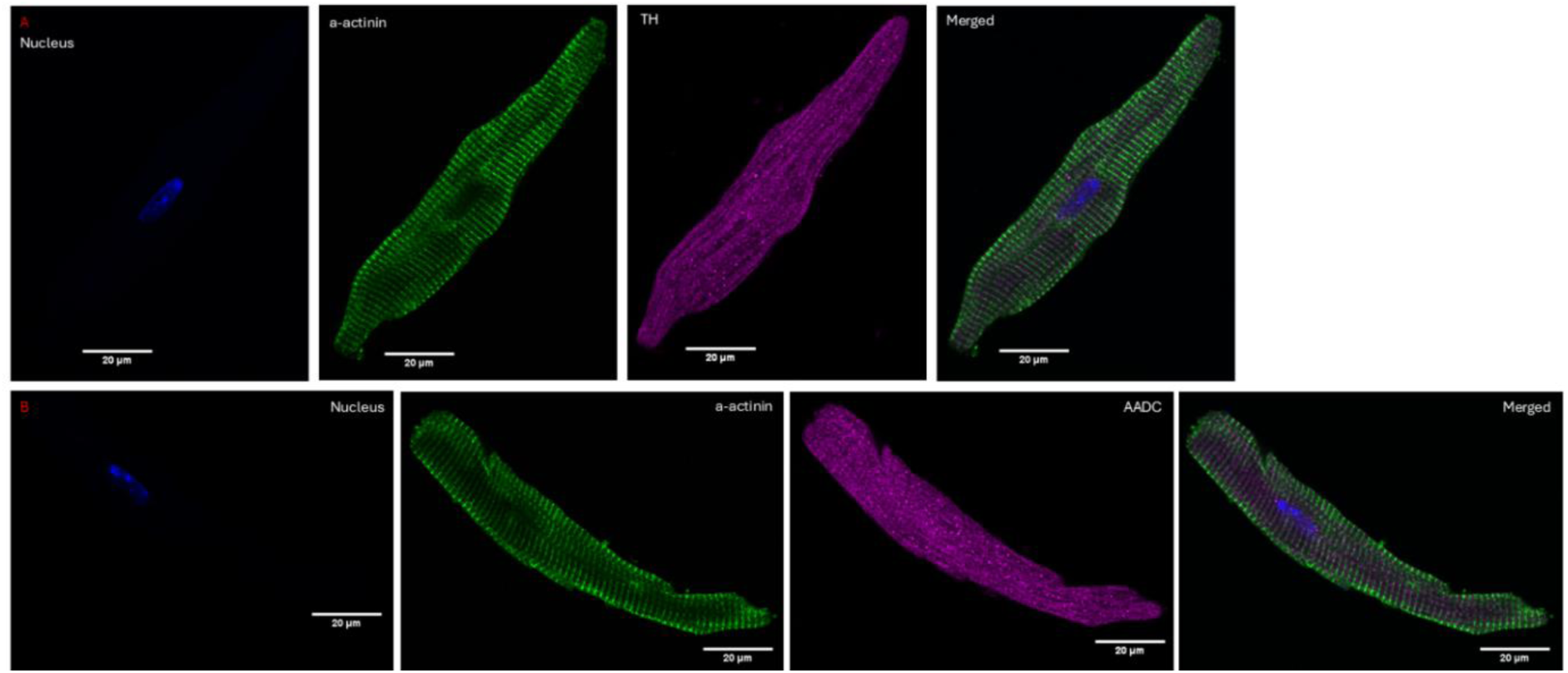
Cytosolic localisation of catecholamine biosynthetic enzymes in cardiomyocytes. Representative immunofluorescence images of isolated CD1 mouse ventricular cardiomyocytes stained for α-actinin (green), nuclei (blue), and tyrosine hydroxylase (TH, magenta; A) or aromatic L-amino acid decarboxylase (AADC, magenta; A). Merged images are shown in the right panels. Both TH and AADC exhibit predominantly cytosolic distribution with diffuse intracellular staining. Scale bars: 20 µm.

### 3.2 Cardiomyocyte-like cells exhibit functional catecholaminergic vesicular uptake and storage

H9C2 cells showed clear uptake and intracellular accumulation of both FFN102 and FFN270, indicating the presence of functional monoamine uptake and vesicular storage activity. FFN102 and FFN270 produced distinct intracellular fluorescence patterns, consistent with the ability of these cardiomyocyte-like cells to accumulate dopamine- and norepinephrine-associated fluorescent substrates, respectively (Figure 4). Treatment with reserpine reduced FFN fluorescence signal, supporting the involvement of vesicular monoamine transport in intracellular dye retention. Together, these findings provide functional evidence that H9C2 cells possess catecholaminergic uptake capacity and VMAT-dependent vesicular storage mechanisms.

**Figure 4.**
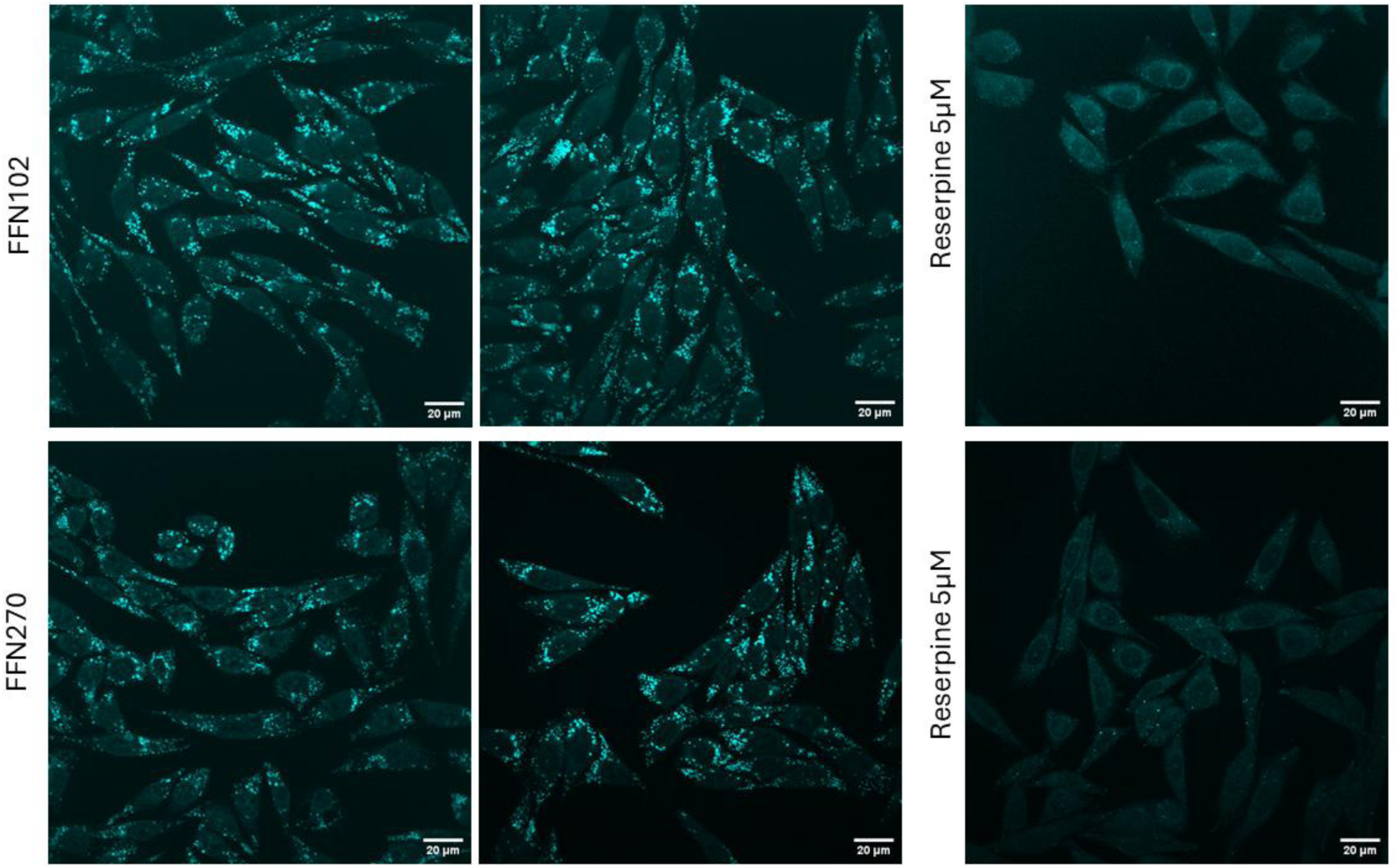
Functional catecholaminergic vesicle uptake in H9C2 cells. Representative fluorescence images of H9C2 cells following incubation with fluorescent false neurotransmitters FFN102 (top row) and FFN270 (bottom row). Both compounds demonstrate intracellular accumulation consistent with monoamine transporter-mediated uptake and vesicular storage. Scale bar: 20μm.

### 3.3 Ischaemia–reperfusion injury induces coordinated upregulation of catecholamine biosynthesis and vesicular handling in the myocardium

Masson’s trichrome staining confirmed successful induction of LAD infarction. I/R hearts exhibited prominent collagen-rich fibrotic deposition consistent with infarct formation, while sham-operated hearts showed preserved myocardial architecture with no evidence of fibrosis (Figure 5).

**Figure 5.**
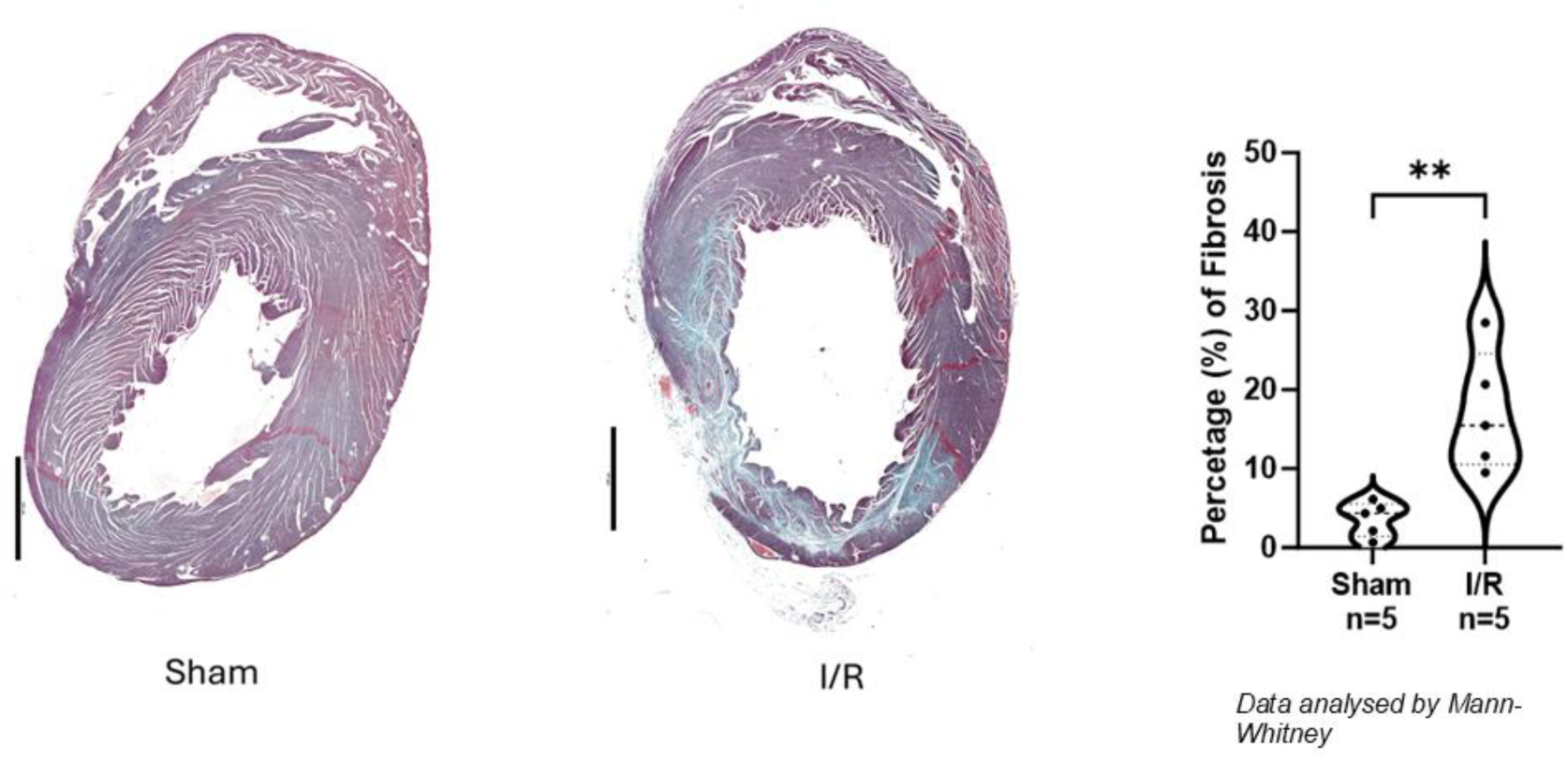
Histological confirmation of infarction following ischaemia–reperfusion injury. Representative Masson’s trichrome staining of rat ventricular tissue from sham-operated and ischaemia–reperfusion (I/R) hearts. Fibrotic tissue deposition is evident in I/R hearts, consistent with infarct formation, whereas sham hearts show no evidence of fibrosis. Scale bar 200μm.

In sham hearts, TH, AADC and DBH were detected in cardiomyocytes (Immunoreactivity score [IRS] 2.2 ±0.84, 5.8 ±2.05, 6 ±1.58, respectively).

Following I/R injury, a significant increase in both the proportion of positively stained cardiomyocytes and IRS scores was observed for all three enzymes (TH: p = 0.008; AADC: p = 0.031; DBH: p = 0.008), indicating coordinated upregulation of the catecholamine biosynthetic pathway in response to injury (Figure 6).

**Figure 6.**
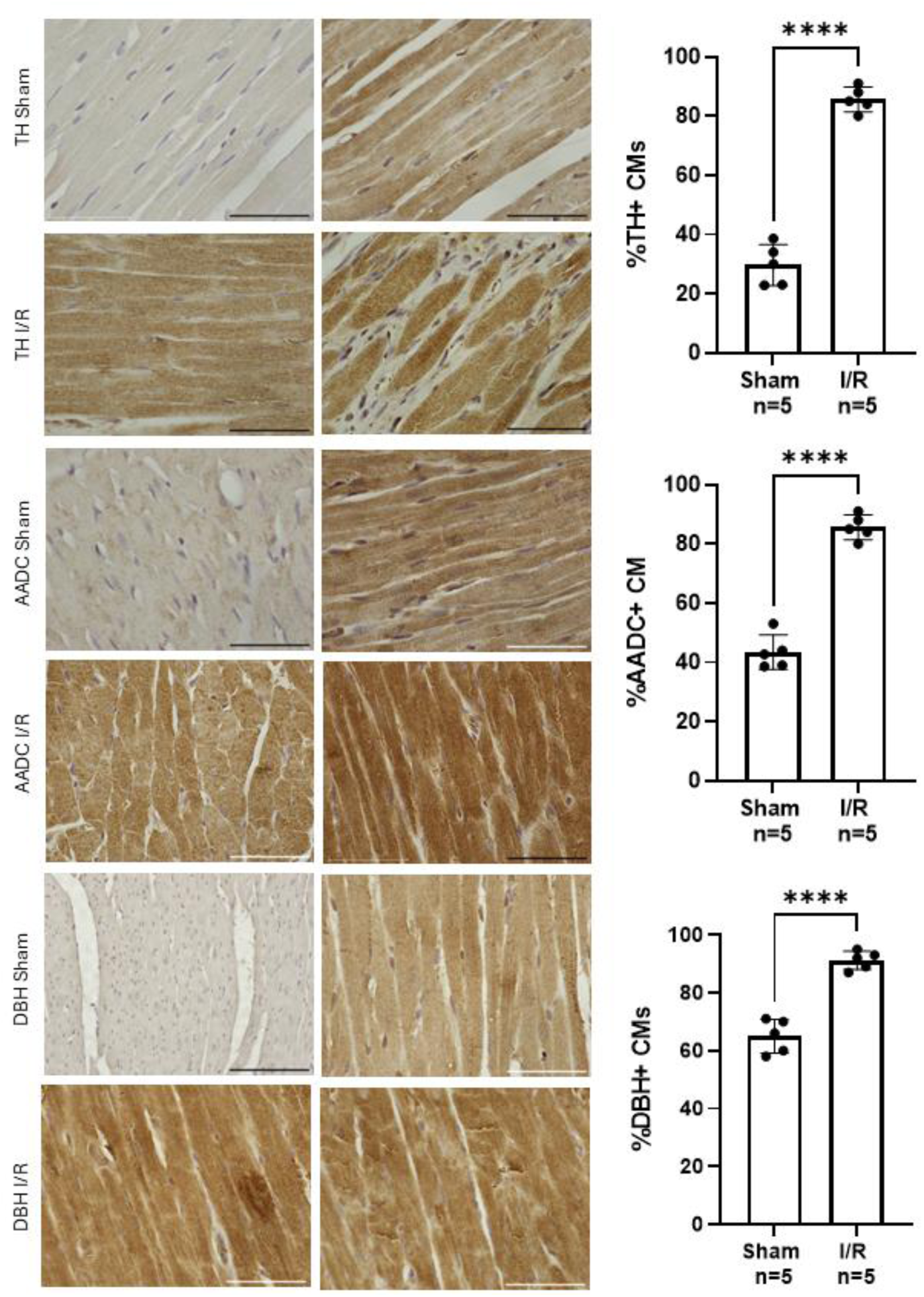
Ischaemia–reperfusion induces upregulation of catecholamine biosynthetic enzymes in cardiomyocytes. Representative immunohistochemical staining of rat ventricular tissue from sham and I/R hearts for TH, AADC and DBH. Quantification shows the percentage of positive cardiomyocytes for each enzyme. Data are presented as mean ± SD (n = 5 per group). Statistical comparisons were performed using unpaired t-test for % of positive cells/ Mann–Whitney test for IRS score. *p < 0.05, **p < 0.01, ***p < 0.001, ****p < 0.0001. Scale bar 50μm.

To further evaluate catecholamine storage and handling capacity, vesicular monoamine transporter expression was assessed. VMAT1 expression was low in sham hearts and did not significantly change following I/R injury (p = 0.119). In contrast, VMAT2 expression was markedly increased in I/R hearts compared to sham controls (p = 0.008), indicating an enhanced capacity for vesicular monoamine uptake and storage in the injured myocardium.

Monoamine oxidase A (MAO-A), a key enzyme involved in catecholamine degradation, was abundantly expressed in sham hearts, supporting the presence of intrinsic catecholamine metabolic capacity within cardiomyocytes (Figure 7).

**Figure 7.**
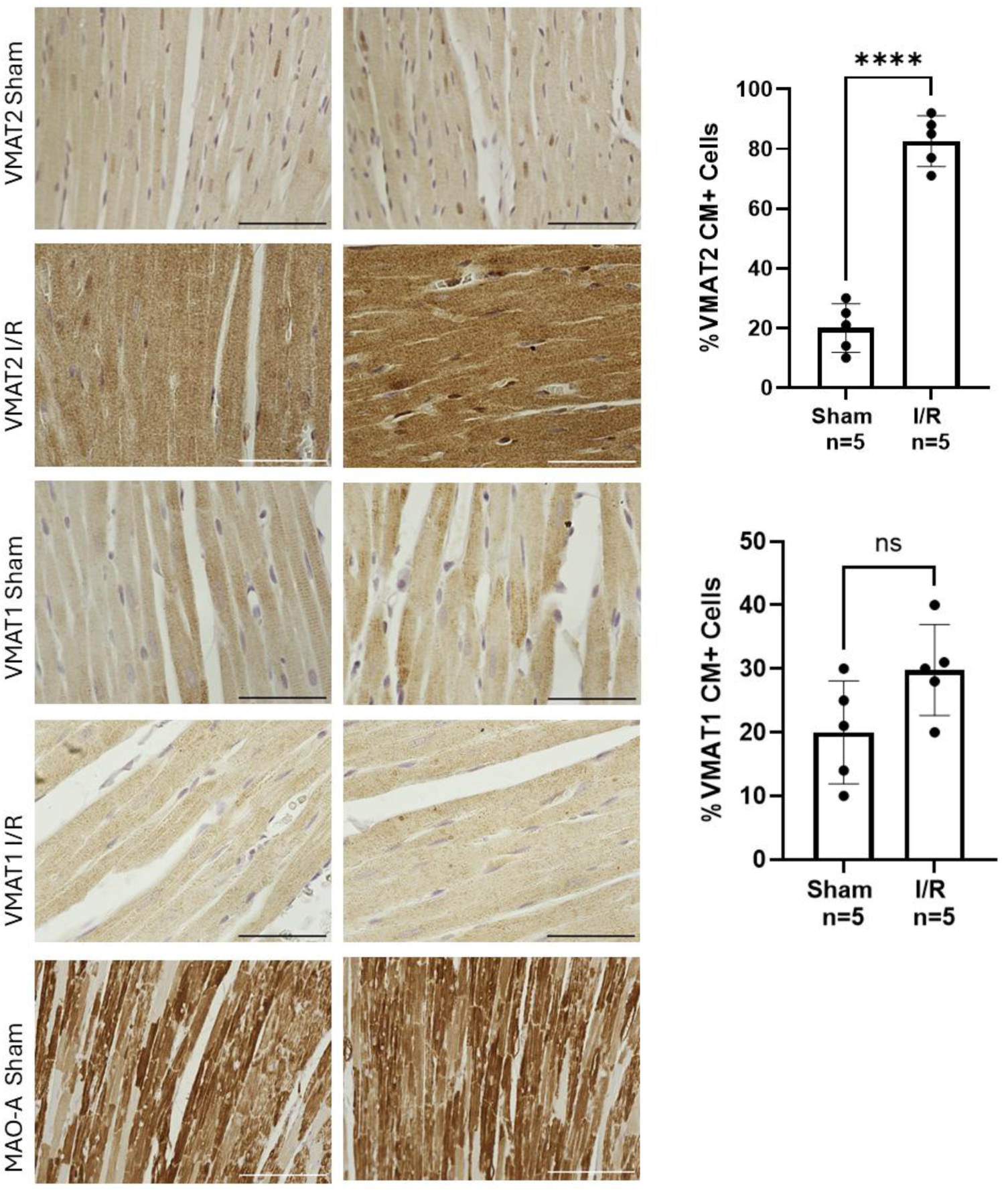
Differential regulation of catecholamine storage and degradation pathways following I/R injury. Representative immunohistochemical staining of rat ventricular tissue from sham and I/R hearts for vesicular monoamine transporter 1 (VMAT1), vesicular monoamine transporter 2 (VMAT2), and monoamine oxidase A (MAO-A). Quantification of VMAT1 and VMAT2 expression is shown as percentage of positive cardiomyocytes. Data are presented as mean ± SEM (n = 5 per group). Statistical comparisons were performed using unpaired t-test for % of positive cells/ Mann–Whitney test for IRS score. **p < 0.01, ****p < 0.0001; ns, not significant. Scale bar: 50μm.

### 3.4 Inhibition of catecholamine biosynthesis impairs viability and disrupts organelle homeostasis

Pharmacological inhibition of TH, AADC and DBH resulted in a dose-dependent increase of H9C2 dead cells, which was further exacerbated following H₂O₂-induced oxidative stress (figure 8). In keeping with these results, catecholamine synthesis enzyme inhibition reduced the metabolic activity of the cells (figure 9, p<0.0001) while increased the activity of caspase 3 / 7.

**Figure 8.**
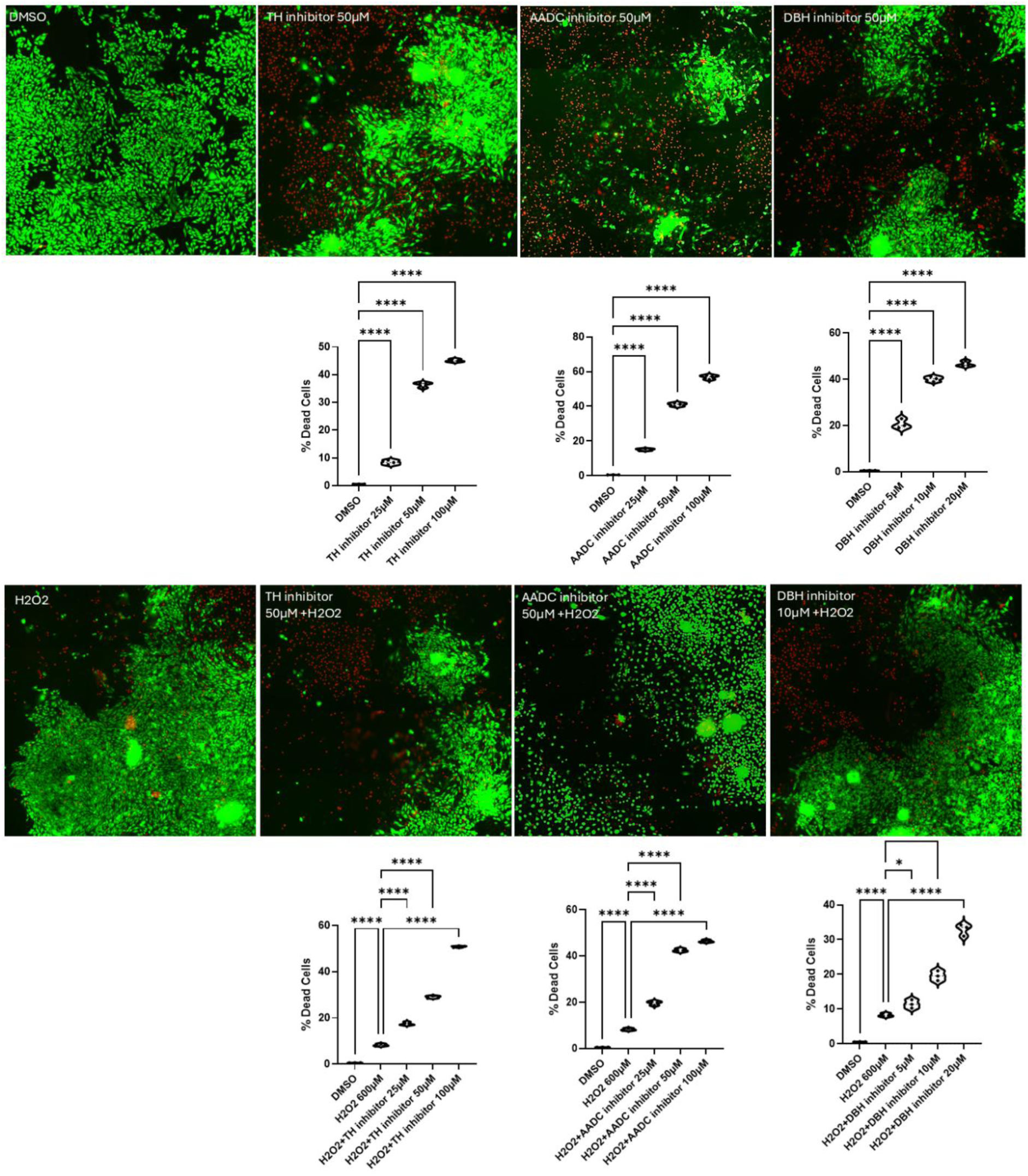
Inhibition of catecholamine biosynthesis increases cell death and reduces stress tolerance in H9C2 cells. Representative live/dead fluorescence images of H9C2 cells treated with inhibitors of TH, AADC and DBH under basal conditions (top row) and following H₂O₂ exposure (bottom row). Live cells are shown in green and dead cells in red (n = 3 independent experiments). Statistical comparisons were performed using ANOVA. ****p < 0.0001.

**Figure 9.**
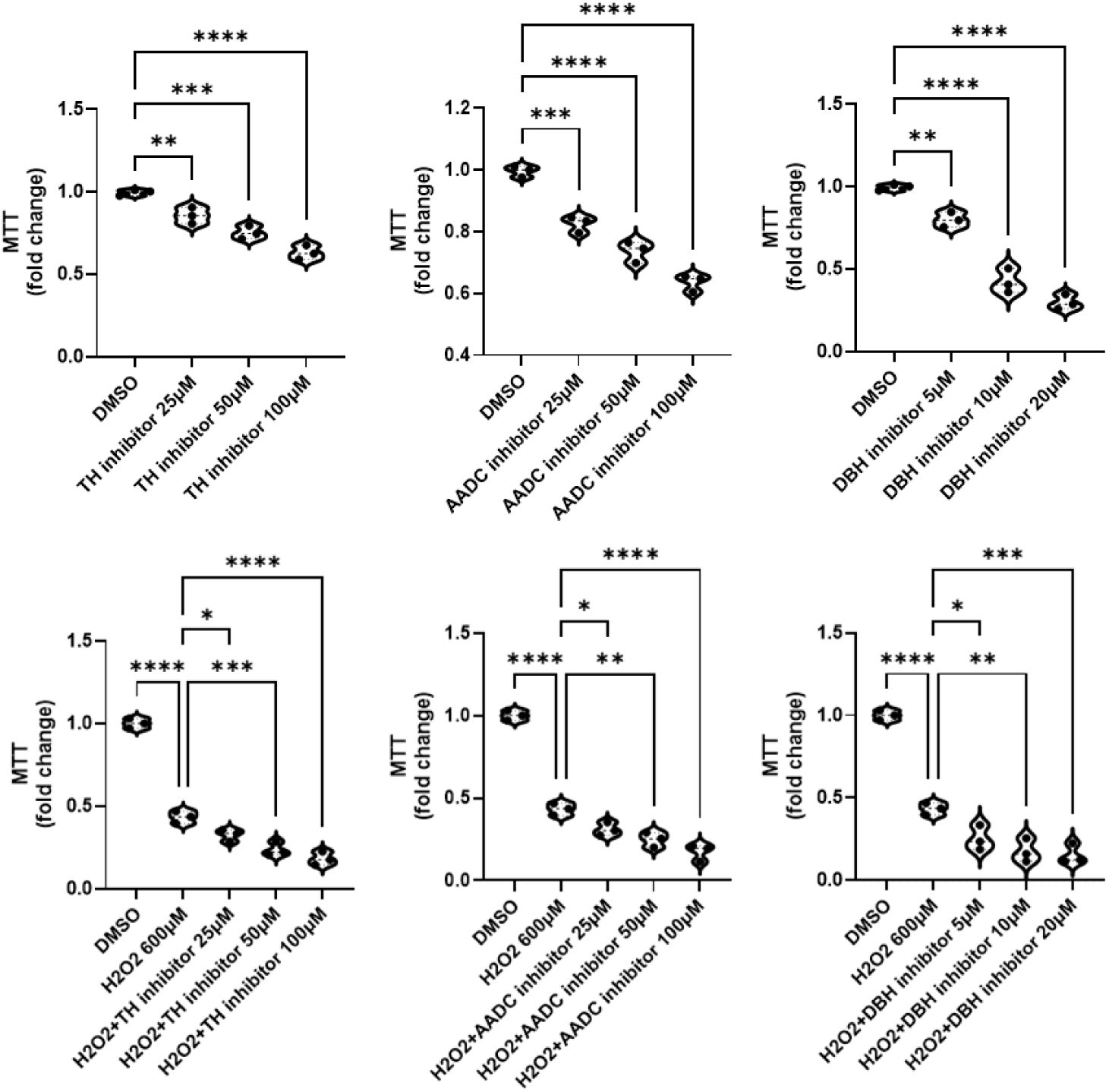
Inhibition of catecholamine biosynthesis reduces metabolic activity in H9C2 cells at baseline and following oxidative stress. MTT assay demonstrating dose-dependent reduction in metabolically active cells following inhibition of TH, AADC, and DBH under basal conditions (top panels) and after H₂O₂ exposure (bottom panels). Data are presented as fold change relative to DMSO control (n = 3 independent experiments). Statistical comparisons were performed using ANOVA. *p < 0.05, **p < 0.01, ***p < 0.001, ****p < 0.0001.

Intracellular and mitochondrial ROS level was significantly increased after blocking the catecholamine synthesis pathway both in baseline conditions and in under H₂O₂-induced oxidative stress. Mitochondrial membrane depolarisation was observed supporting mitochondria dysfunction leading to apoptosis (figure 10).

**Figure 10.**
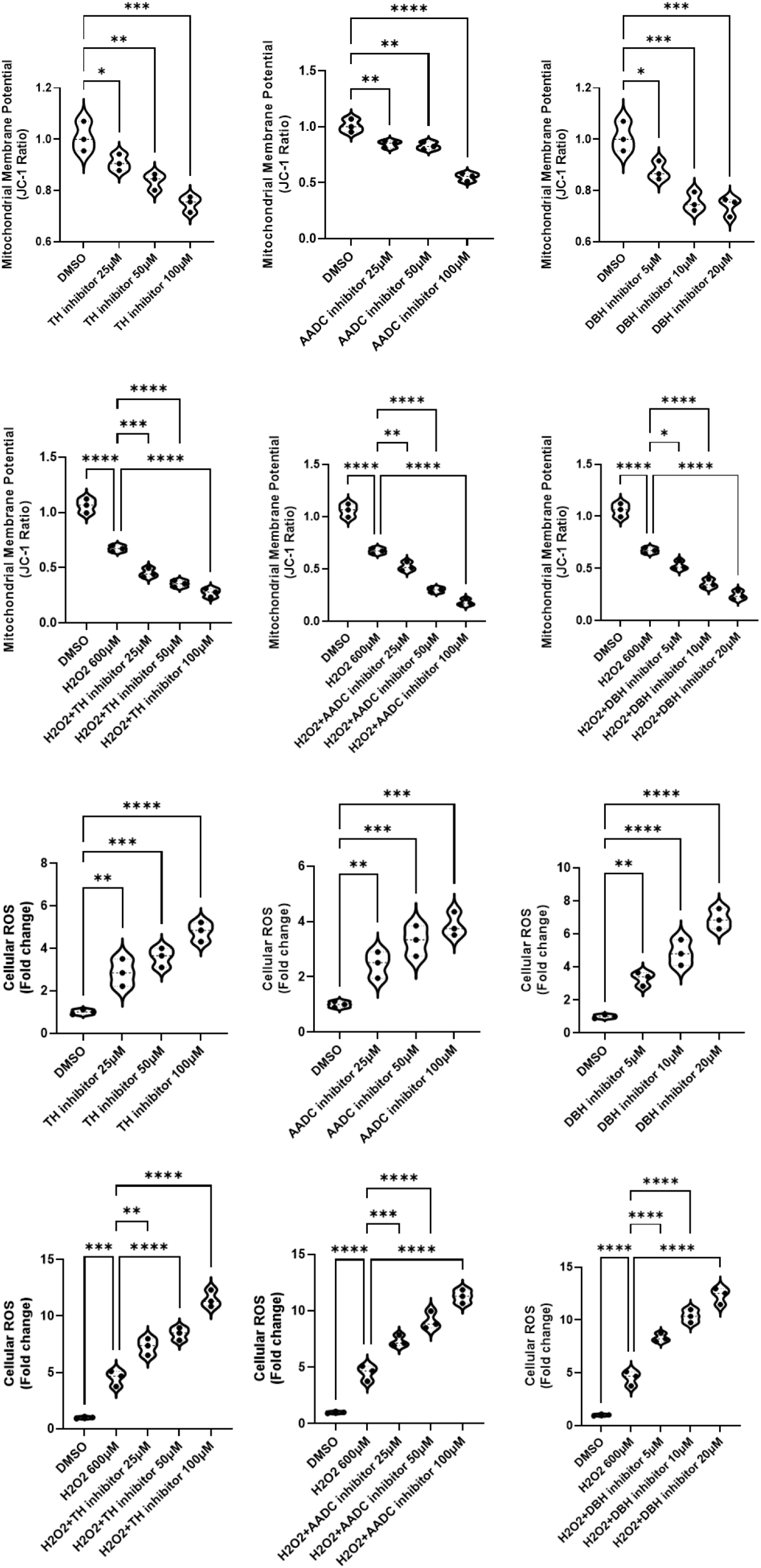
Effects of TH, AADC, and DBH inhibition on mitochondrial membrane potential and cellular ROS production under basal and oxidative stress conditions. Violin plots show mitochondrial membrane potential (ΔΨm), assessed by the JC-1 red/green fluorescence ratio, and intracellular reactive oxygen species (ROS) levels expressed as fold change relative to DMSO-treated controls. Under basal conditions (top and third rows), treatment with tyrosine hydroxylase (TH), aromatic L-amino acid decarboxylase (AADC), or dopamine β-hydroxylase (DBH) inhibitors resulted in a dose-dependent reduction in mitochondrial membrane potential and a concomitant increase in cellular ROS levels compared with DMSO controls. Under oxidative stress induced by H₂O₂ (600 μM) (second and fourth rows), co-treatment with TH, AADC, or DBH inhibitors further exacerbated mitochondrial depolarization and significantly increased ROS generation in a concentration-dependent manner relative to H₂O₂ alone. Each violin plot represents the distribution of individual biological replicates, with central lines indicating median values and dotted lines indicating quartiles. Statistical significance was determined using one-way ANOVA followed by multiple-comparison testing. *P < 0.05, **P < 0.01, ***P < 0.001, ***P < 0.0001.

Lysosomal and endoplasmic reticulum organelles were increased in number and size with increased activity to suggest that multiple organellar stress pathways contribute to cell damage and death (figure 11).

**Figure 11.**
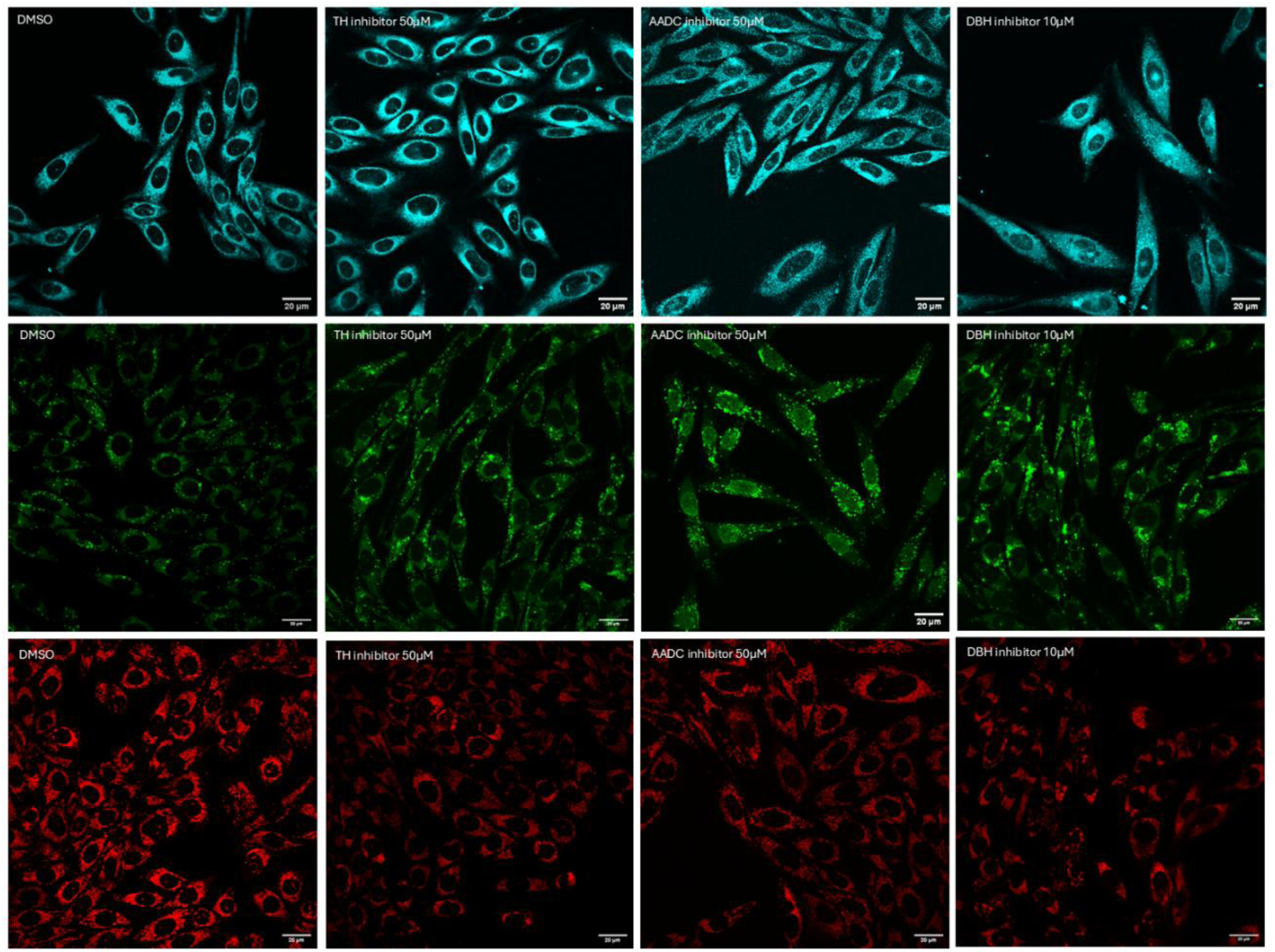
Pharmacological inhibition of catecholamine biosynthesis alters organelle homeostasis in H9C2 cardiomyoblasts. Representative confocal fluorescence images of H9C2 cells following treatment with vehicle control (DMSO), tyrosine hydroxylase (TH) inhibitor (50 μM), aromatic L-amino acid decarboxylase (AADC) inhibitor (50 μM), or dopamine β-hydroxylase (DBH) inhibitor (10 μM). The upper panel shows endoplasmic reticulum staining, the middle panel lysosomal staining, and the lower panel mitochondrial staining. Images are representative of independent biological experiments performed under identical imaging conditions. Scale bar = 20 μm.

The results were similar for all three compounds suggesting that all parts of the catecholamine synthesis pathway are essential.

These findings suggest that endogenous catecholamine biosynthesis plays a critical central role in maintaining cardiomyocyte viability, homeostasis and stress resilience.

### 3.5 Catecholamine biosynthesis inhibition reprograms H9c2 metabolic signalling toward an energy-conserving state

Under basal conditions, inhibition of TH, AADC and DBH was associated with marked suppression of mTOR signalling. This was demonstrated by a significant reduction in the p-mTOR/mTOR ratio, together with decreased phosphorylation of the downstream mTORC1 targets S6 ribosomal protein and 4EBP1. These findings indicate that disruption of catecholamine biosynthesis reduces mTORC1 activity and attenuates downstream protein-synthesis-associated signalling.

A similar pattern was observed under oxidative stress conditions. Exposure to H₂O₂ reduced mTOR pathway activity, and combined treatment with H₂O₂ and catecholamine-biosynthesis inhibitors further enhanced suppression of p-mTOR, p-S6 ribosomal protein and p-4EBP1.

This suggests that inhibition of catecholamine biosynthesis sensitises H9c2 cells to oxidative-stress-associated repression of anabolic signalling.

In parallel, catecholamine-biosynthesis inhibition was associated with activation of energy-stress signalling. Increased phosphorylation of LKB1 and ACC was observed, consistent with activation of an LKB1–AMPK-related metabolic stress response and inhibition of energy-consuming lipid biosynthetic pathways. Phosphorylation of ULK1 was also increased, indicating engagement of autophagy-related signalling pathways. These changes were evident both at baseline and during H₂O₂ exposure, suggesting that catecholamine-biosynthesis inhibition shifts H9c2 cells towards an energy-conserving, stress-adaptive phenotype (Figure 12).

**Figure 12.**
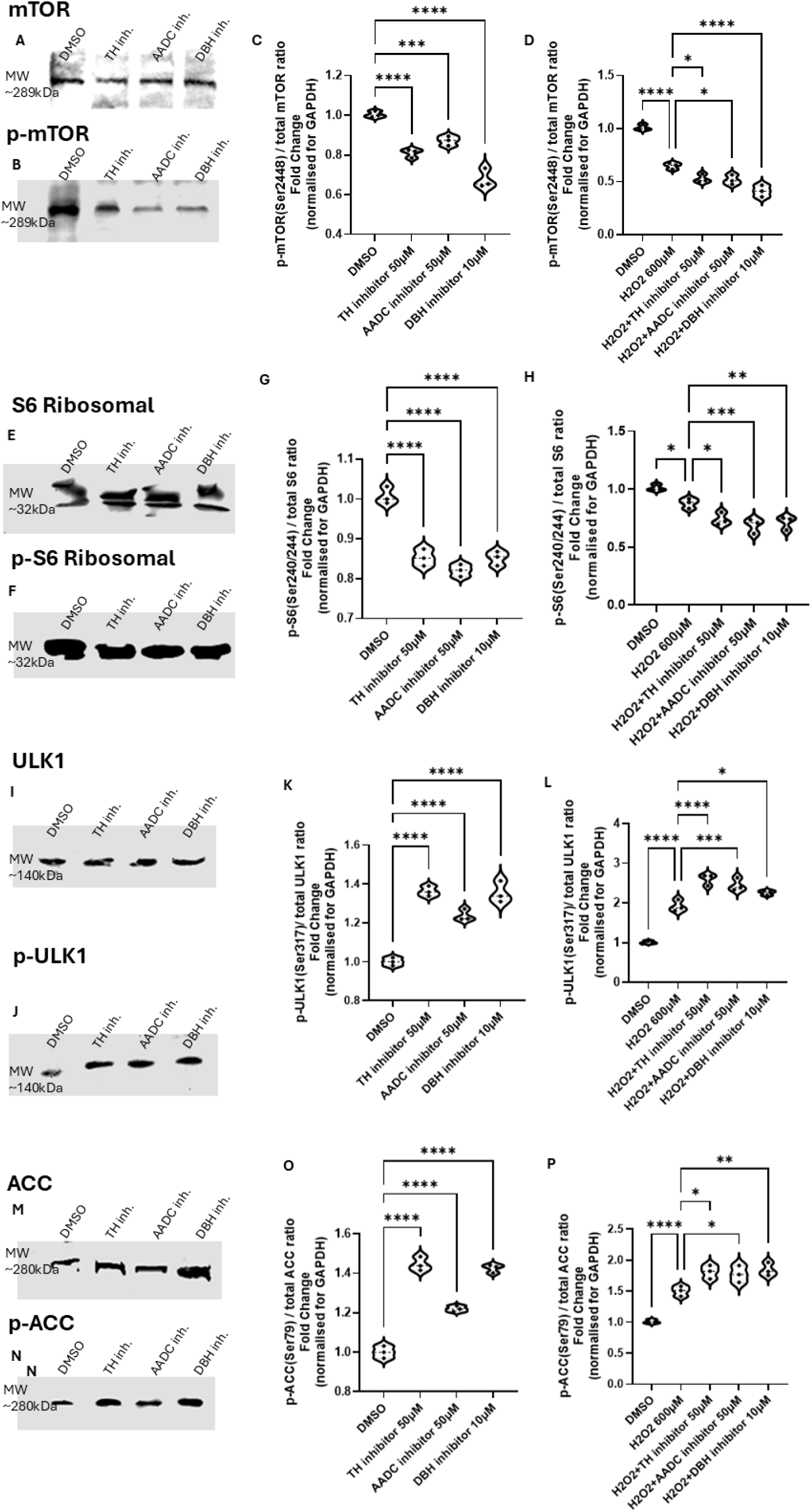
Inhibition of catecholamine biosynthesis suppresses mTORC1 signalling and activates energy stress pathways. Representative western blots and quantification showing the effects of TH, AADC and DBH inhibition under basal conditions and following H₂O₂-induced oxidative stress in H9c2 cells. (A, B) Representative western blots showing total mTOR and phosphorylated mTOR expression. (C, D) Quantification of the p-mTOR/mTOR ratio under basal conditions and following H₂O₂ exposure. (E, F) Representative western blots showing total S6 ribosomal protein and phosphorylated S6 ribosomal protein expression. (G, H) Quantification of the p-S6/S6 ratio. (I, J) Representative western blots showing total ULK1 and phosphorylated ULK1 expression. (K, L) Quantification of the p-ULK1/ULK1 ratio. (M, N) Representative western blots showing total ACC and phosphorylated ACC expression. (O, P) Quantification of the p-ACC/ACC ratio. Data demonstrate that inhibition of catecholamine biosynthesis suppresses mTORC1 signalling, as shown by reduced phosphorylation of mTOR and its downstream targets S6 ribosomal protein and 4EBP1, while increasing phosphorylation of energy-stress and autophagy-related markers, including ULK1 and ACC. Statistical analysis was performed using one-way ANOVA. *p < 0.05, **p < 0.01, ***p < 0.001, ****p < 0.0001.

Collectively, these data demonstrate that inhibition of TH, AADC and DBH suppresses mTORC1-dependent anabolic signalling while promoting activation of energy-sensing and autophagy-related pathways. This metabolic reprogramming is characterised by reduced phosphorylation of mTOR, S6 ribosomal protein and 4EBP1, together with increased phosphorylation of LKB1, acetyl-CoA carboxylase (ACC) and ULK1. These findings support the concept that disruption of catecholamine biosynthesis compromises cellular bioenergetic homeostasis and may contribute to impaired H9c2 cell survival, particularly under oxidative stress conditions.

### 3.5 Tyrosine hydroxylase inhibition promotes electrophysiological instability in ex vivo hearts

In Langendorff-perfused mouse hearts, pharmacological inhibition of tyrosine hydroxylase with metyrosine induced ventricular arrhythmias in 5 out of 6 hearts. Ventricular tachycardia (VT) was inducible by burst pacing in 3 hearts, with episodes ranging from short, self-terminating runs to prolonged sustained arrhythmias lasting up to 28 minutes. In one preparation, VT alternated with episodes of ventricular fibrillation (VF), indicating severe electrical instability.

In addition to sustained arrhythmias, non-sustained ventricular tachycardia (NSVT) was observed in 2 hearts, occurring spontaneously and following burst pacing. Both monomorphic and polymorphic VT were observed, including one episode of bidirectional VT (Figure 13).

**Figure 13.**
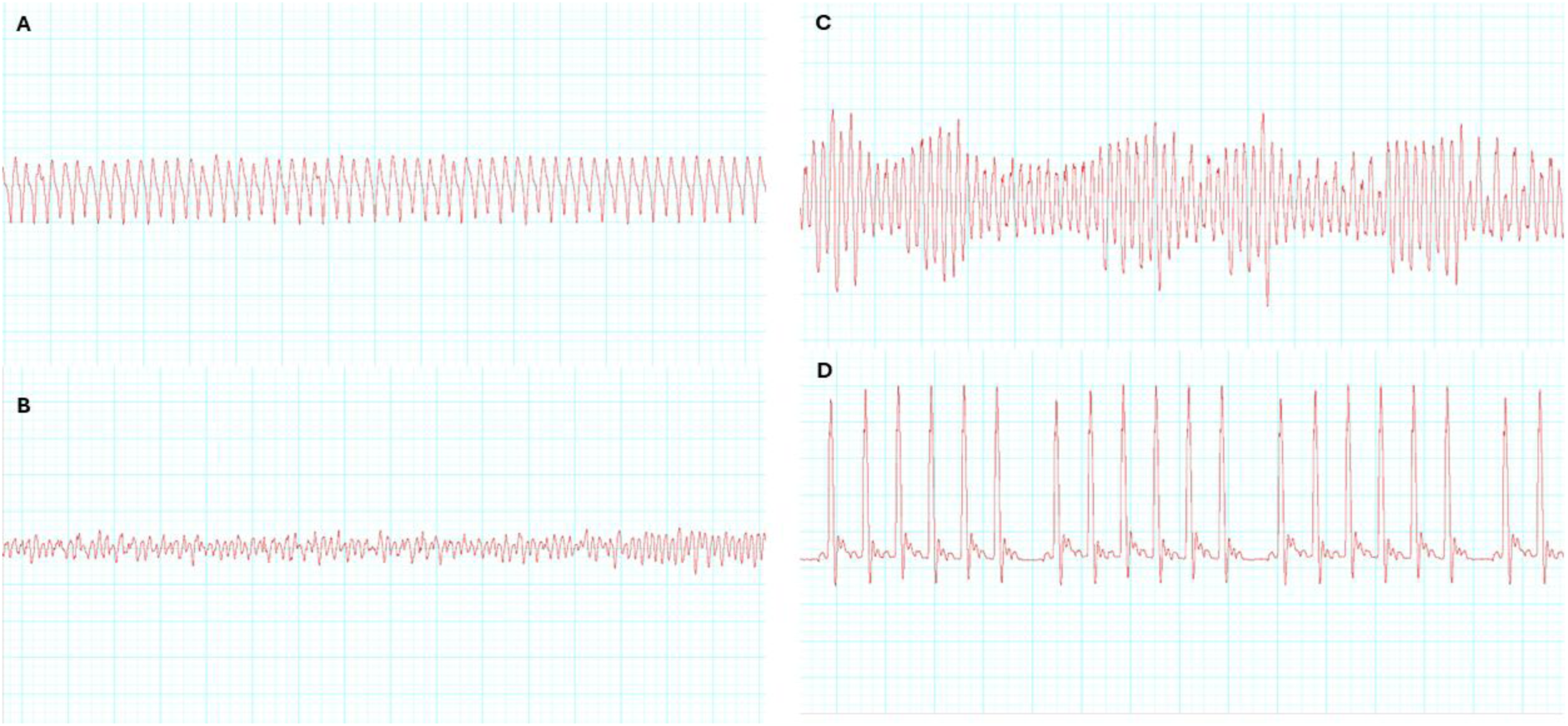
Tyrosine hydroxylase inhibition induces ventricular arrhythmias in ex vivo hearts. Representative electrogram traces from Langendorff-perfused mouse hearts following treatment with the tyrosine hydroxylase inhibitor metyrosine. (A) Sustained monomorphic ventricular tachycardia (VT). (B) Ventricular fibrillation (VF). (C) Polymorphic ventricular tachycardia. Traces illustrate the spectrum of ventricular arrhythmias observed following inhibition of catecholamine biosynthesis. (D) Weckenbach observed following TH inhibition, indicating atrioventricular conduction disturbance.

In the remaining heart, atrioventricular conduction abnormalities were observed, with Wenckebach periodicity developing following metyrosine perfusion.

Importantly, all electrophysiological abnormalities were reversible following washout with Krebs solution, supporting a direct functional effect of catecholamine biosynthesis inhibition on cardiac electrical stability.

## 4. Discussion

Our findings support the existence of a cardiomyocyte-intrinsic catecholaminergic machinery, characterised by coordinated expression of the key catecholamine biosynthetic enzymes TH, AADC, DBH and PNMT, together with vesicular monoamine handling through VMAT1 and VMAT2 (23). Functional interrogation of this pathway demonstrated its relevance at both the cellular and whole-heart levels.

Pharmacological inhibition of catecholamine biosynthesis impaired cardiomyocyte viability, disrupted organelle homeostasis and triggered a metabolic stress response marked by suppression of mTOR signalling and activation of the LKB1–AMPK–ULK1 axis. Although these mechanistic data are based largely on pharmacological inhibition, the concordant effects observed following inhibition of several sequential enzymes in the catecholamine biosynthetic pathway support the biological relevance of this axis. These cellular perturbations were accompanied by pronounced ventricular electrical instability in ex vivo hearts, including induction of ventricular arrhythmias following tyrosine hydroxylase inhibition. Collectively, these findings identify a previously unrecognised intrinsic monoaminergic regulatory machinery within cardiomyocytes that integrates metabolic homeostasis with electrophysiological stability.

Early studies reported measurable tyrosine hydroxylase activity and norepinephrine content in rat cardiac tissue, suggesting that catecholamine biosynthesis may occur outside classical neuroendocrine organs (24). More recently, endogenous release of 6-cyanodopamine has been described in rat hearts and proposed to modulate chronotropism and inotropism, although its cellular origin was not defined (25). Other work has identified monoamine-related enzymes in cardiomyocytes, including AADC and monoamine oxidase A, in the context of cardiomyocyte serotonin synthesis (26), while transcriptomic analyses have suggested DBH mRNA expression in myocardial tissue (27). Together, these studies provided important evidence for discrete components of monoamine biology within the heart, but did not establish a complete, coordinated and functionally active catecholaminergic machinery in cardiomyocytes, or define its significance for myocardial homeostasis and electrical stability (28).

Our findings extend this earlier work by demonstrating coordinated cardiomyocyte expression of catecholamine biosynthetic enzymes together with vesicular monoamine transport machinery. By combining molecular, cellular and whole-heart functional analyses, we provide evidence that this machinery is not merely transcriptionally detectable but biologically active. Inhibition of catecholamine biosynthesis disrupted cardiomyocyte homeostasis and precipitated ventricular electrical instability, supporting a role for this intrinsic pathway in linking monoamine handling to metabolic and electrophysiological regulation. Thus, our study advances the concept of cardiac monoamine biology from isolated enzymatic observations to an integrated cardiomyocyte-intrinsic regulatory machinery.

The marked effects observed following inhibition of catecholamine biosynthesis support a functional role for intrinsic catecholaminergic signalling in the maintenance of cardiomyocyte homeostasis. Pharmacological inhibition of TH, AADC, and DBH impaired cardiomyocyte viability, and induced coordinated cellular stress responses, including mitochondrial depolarisation, lysosomal and endoplasmic reticulum expansion, and activation of apoptotic signalling. Cells were also incubated with H_2_O_2_, known to lead to ischaemia like oxidative stress (29, 30), and the above effects were amplified further, supporting a role for intrinsic catecholaminergic signalling in maintaining cardiomyocyte stress resilience.

Mechanistically, disruption of catecholamine biosynthesis was associated with suppression of mTOR signalling (31) and activation of the LKB1–AMPK–ULK1 axis, consistent with a shift towards an energy-deprived, catabolic cellular state (32-35). These findings suggest that intrinsic catecholaminergic signalling may support cardiomyocyte metabolic homeostasis by preserving organelle integrity and restraining maladaptive stress pathway activation (36-38). Collectively, our data identify this pathway as a previously unrecognised regulator of cardiomyocyte viability, metabolic adaptation and stress tolerance.

Importantly, the cellular consequences of catecholamine biosynthesis inhibition were accompanied by marked electrophysiological instability at the whole-heart level. In Langendorff-perfused hearts, tyrosine hydroxylase inhibition induced ventricular arrhythmias in the majority of preparations, including sustained and polymorphic ventricular tachycardia and ventricular fibrillation. Catecholaminergic signalling is a well-established regulator of cardiac excitability and arrhythmogenesis, principally through sympathetic nervous system activation and β-adrenergic receptor signalling, calcium handling and excitation–contraction coupling (3, 39-42). Our findings extend this paradigm by suggesting that catecholamine-dependent regulation of myocardial electrical stability may also arise, at least in part, from within cardiomyocytes themselves (43). This concept is consistent with emerging evidence that cardiomyocytes harbour intrinsic neurotransmitter-related pathways capable of modulating cardiac function independently of classical neurohumoral input. Together, these findings support a previously underappreciated role for cardiomyocyte-intrinsic catecholaminergic signalling in linking cellular metabolic stress to ventricular electrical instability.

Our findings are consistent with a growing body of evidence indicating that cardiomyocytes possess intrinsic neurotransmitter regulatory systems in addition to responding to classical neurohumoral input. Catecholaminergic signalling is a well-established determinant of cardiac excitability, metabolism and stress adaptation through sympathetic activation and β-adrenergic pathways. However, prior reports of tyrosine hydroxylase activity, norepinephrine content or isolated catecholamine-related enzymes within the myocardium did not establish a coordinated, functional cardiomyocyte-intrinsic catecholaminergic machinery. More recently, the identification of an endogenous cholinergic signalling pathway within cardiomyocytes has challenged the long-held view that these cells are solely passive targets of neuronal regulation (15, 44). In this context, our data extend the concept of intrinsic cardiac neurotransmitter signalling by supporting a cardiomyocyte-intrinsic catecholaminergic pathway that contributes to metabolic homeostasis and electrophysiological stability. The associated suppression of mTOR signalling and activation of LKB1–AMPK–ULK1 pathways further aligns with established links between metabolic stress responses, autophagy and cardiomyocyte survival (45).

Several limitations should be considered. First, many functional experiments were performed in H9C2 cells, which, although widely used to study cardiomyocyte stress responses, do not fully recapitulate the structural, metabolic or electrophysiological phenotype of mature cardiomyocytes (46). However, complementary findings in isolated adult cardiomyocytes, hiPSC-derived cardiomyocytes and Langendorff-perfused hearts support the physiological relevance of the pathway beyond this model. Second, a substantial component of the mechanistic evidence relies on pharmacological inhibition of catecholamine biosynthesis.

Although the concordant effects observed following inhibition of TH, AADC and DBH argue against a single compound-specific effect, off-target actions cannot be fully excluded, and future genetic or other orthogonal approaches will be required to confirm pathway specificity. Third, this study did not directly quantify intracellular or secreted catecholamines, and direct measurements of dopamine, norepinephrine, epinephrine and related intermediates will be important to define the biochemical output and regulation of this pathway. Finally, while tyrosine hydroxylase inhibition was associated with impaired electrophysiological stability, the precise mechanisms linking impaired catecholamine biosynthesis to ventricular arrhythmia susceptibility remain to be defined, including the relative contributions of metabolic stress signalling, mitochondrial dysfunction, calcium handling and ion-channel regulation.

In conclusion, our findings support the existence of an intrinsic catecholaminergic machinery within cardiomyocytes that contributes to the regulation of metabolic homeostasis and electrical stability. Disruption of this pathway impaired cellular stress responses, activated energy-sensing signalling pathways and increased susceptibility to ventricular arrhythmias. Together, these findings provide new insight into cardiomyocyte-intrinsic monoaminergic signalling and identify a previously unrecognised mechanism linking metabolic stress responses to myocardial electrophysiological regulation.

## Acknowledgments

The authors thank the Microscopy Facility, Dunn School of Pathology, University of Oxford, for support with microscopy.

